# Genetic diversity, recombination and cross-species transmission of a waterbird gammacoronavirus in the wild

**DOI:** 10.1101/2023.06.08.544213

**Authors:** Sarah François, Salik Nazki, Stephen H. Vickers, Guillaume Fournié, Christopher M. Perrins, Andrew J. Broadbent, Oliver G. Pybus, Sarah C. Hill

## Abstract

Viruses emerging from wildlife can cause outbreaks in humans and domesticated animals. Predicting the emergence of future pathogens and mitigating their impacts requires an understanding of what shapes virus diversity and dynamics in wildlife reservoirs. In order to better understand coronavirus ecology in wild species, we sampled birds within a coastal freshwater lagoon habitat across five years, focussing on a large population of mute swans (*Cygnus olor*) and the diverse species that they interact with. We discovered and characterised the full genome of a divergent gammacoronavirus belonging to the *Goose coronavirus CB17* species. We investigated the genetic diversity and dynamics of this gammacoronavirus using untargeted metagenomic sequencing of 223 fecal samples from swans of known age and sex, and RT-PCR screening of 1632 additional bird samples. The virus circulated persistently within the bird community; virus prevalence in mute swans exhibited seasonal variations, but did not change with swan age-class or epidemiological year. One whole genome was fully characterised, and revealed that the virus originated from a recombination event involving an undescribed gammacoronavirus species. Multiple lineages of this gammacoronavirus co-circulated within our study population. Viruses from this species have recently been detected in aquatic birds from both the Anatidae and Rallidae families, implying that host species habitat sharing may be important in shaping virus host range. As the host range of the *Goose coronavirus CB17* species is not limited to geese, we propose that this species name should be updated to “*Waterbird gammacoronavirus 1*”. Non-invasive sampling of bird coronaviruses may provide a tractable model system for understanding the evolutionary and cross-species dynamics of coronaviruses.

## 3. Introduction

Viruses emerging from wildlife cause major outbreaks in humans and domesticated animals. The rate of pathogen spillover from wild mammals and birds is increasing, with globalisation and anthropogenic land-use change leading to increasing contact between humans and wildlife (1–3). Viruses are of particular public health concern and represent 75% of new human pathogens discovered since 1980 (4,5). The threats posed by emerging coronaviruses emphasise the need to identify and understand viruses with spillover potential in their wildlife reservoirs.

The development and adoption of high-throughput sequencing approaches for the untargeted exploration of virus genomic diversity (i.e. virus metagenomics) has in recent years enabled virus discovery at an unprecedented scale (6,7). Individual-level host information (e.g., age, sex) is often unknown for wild individuals, and wildlife can be hard to capture and sample. Thus most virus metagenomic studies remain descriptive and focussed on cataloguing the viruses found in natural populations (8,9). However, understanding the factors that determine viral diversity and dynamics within wild animal reservoirs is important if we are to predict future outbreaks and mitigate their risks to endangered wildlife, domestic animals, and humans (10,11).

Coronaviruses (family *Coronaviridae*, subfamily *Orthocoronavirinae*) are positive-sense single-stranded RNA viruses that exhibit high evolutionary potential due to their high mutation rates (12,13), propensity to recombine, and long genomes (27-32 kb) compared to most other RNA viruses. This potentially facilitates their adaptation to new hosts (14,15).

Three zoonotic coronaviruses have emerged from wildlife reservoirs in the last twenty years, each posing a substantial threat to global public health: SARS, MERS, and SARS-CoV-2 (16–18). Coronaviruses infect humans and diverse wild and domestic mammals and birds, causing mild or severe respiratory, hepatic, enteric and neurological diseases. Research on coronavirus discovery and characterisation in wildlife continues to focus on bats, because bats are an important reservoir of mammal-infecting alphacoronaviruses and betacoronaviruses, and the latter genus contains zoonotic coronaviruses (19,20). Wild birds are a reservoir of most gammacoronaviruses and deltacoronaviruses (21). Indeed, the first coronavirus to be discovered was a bird-infecting gammacoronavirus, infectious bronchitis virus (IBV). While none of the coronaviruses found in wild birds are known to be zoonotic, IBV likely originated in wild birds (22,23), but is now the causative agent of avian infectious bronchitis, an acute and highly contagious respiratory disease of chickens that causes huge economic losses for the poultry industry (24).

Many wild waterbird species connect habitats across the world through annual migration and are reservoirs of zoonotic and poultry pathogens, particularly avian influenza viruses (25). Despite their possible importance in harbouring and disseminating viruses, knowledge of coronavirus ecology in wild birds is very limited compared to that of coronaviruses in poultry and mammals. Indeed, 11 coronavirus species are currently described in birds whilst 35 are described in mammals. (26). Yet, the broad host range exhibited by some coronaviruses infecting multiple bird species that share aquatic habitats, such as ducks, gulls and shorebirds (27), suggests that the ecology of these viruses is complex. In order to better understand the ecology of coronaviruses in the wild, we conducted repeated cross-sectional sampling in a freshwater coastal lagoon across five years, focussing on a large population of mute swans (*Cygnus olor*) and the birds that they interact with. We investigated the genetic diversity and dynamics of a divergent gammacoronavirus using untargeted metagenomic sequencing of 223 fecal samples from swans and RT-PCR screening of 1632 additional bird samples.

## 4. Methods

### Study community and sampling

Our study site in southern England (50.65378°N, 2.60288°W) harbours a colony of wild mute swans that has been studied since the 1960s. The population comprises approximately 600 to 1000 birds that mix freely with birds of other species. Swan population numbers are higher in the early summer following hatching of cygnets, and in the mid-summer as birds immigrate from local areas into the lagoon for moulting. The swans are ringed and detailed information (age, sex, mating data, parentage) is available for most birds (28,29).

Faecal samples were collected from mute swans on 26 occasions between April 2015 and September 2020, with each visit lasting 2-3 days. Fecal samples from other birds were collected on a subset of these occasions, specifically between July 2019 and September 2020. Mute swans were observed when on land, and non-invasive samples were taken from birds seen to defecate. Almost all swans in this colony are ringed, and each swan’s leg colour-ring (∼5cm long and engraved with a unique three-character and colour code) is large enough to be observed and recorded from a distance without disturbing the bird. When collecting faecal samples from other bird species, we could not get close enough to identify from which species they originated. However, due to their smaller size and our familiarity with swan faeces, we are confident that these samples were not from swans. Approximately 0.5 mL of faeces was collected with sterile plastic spatula into a 1.5 mL tube containing 1 mL of Universal Transport Media (Sterilin) or RNAlater (Thermo Fisher Scientific). The tubes were shaken vigorously and kept on ice or dry ice in the field for up to an hour. Subsequent sample storage and transport was conducted on dry ice, and samples were stored at -80 °C until processing. None of the sampled, ringed mute swans are known to have died within 7 weeks after sampling, and none of the birds had symptoms of disease at the time of sampling based on casual observation from a distance. In total, 1080 samples from mute swans and 671 samples from other bird species were processed. A list of the samples and of their associated metadata is provided in **supplementary table 1**.

### Metagenome acquisition and coronavirus genome reconstruction

Metagenomic sequencing of 223/1080 mute swan fecal samples was conducted as part of a previous study (30). In brief, encapsidated viral nucleic acids were enriched by centrifugation and by DNase treatment. The Boom method was used for extraction of DNA and RNA nucleic acids (31). Reverse transcription was performed using non-ribosomal random hexamers (32). Second strand DNA synthesis was performed with Klenow fragment DNA polymerase. Nucleic acids were purified by phenol/chloroform extraction and ethanol precipitation (33). Each sample was barcoded and paired-end library preparation was performed using an Illumina Nextera XT kit, before sequencing in three multiplexes on an Illumina HiSeq 4000 to generate 150 bp paired-end reads (30). Illumina adaptors were removed and reads were filtered for quality (q30 quality and read length >45nt) using cutadapt 1.18 (34). Reads were assembled de novo into contigs using SPAdes 3.12.0 (k-mer lengths 21, 33, 55, 77) (35). Taxonomic assignment was achieved on contigs of length >900nt through searches against a custom nidovirales protein database that reflected the known diversity within the *Nidovirales* order, using DIAMOND 0.9.22 with an e-value cutoff of <10-5 (36). The nidovirales protein database contained all nidovirus (NCBI:txid76804) proteins deposited in GenBank by 1^st^ September 2020. All contigs that matched nidovirus sequences were used as queries to perform reciprocal searches on the NCBI non-redundant protein sequence database with an e-value cut-off of <10^−3^ to eliminate false positives (37). Coronavirus contig completion and coverage was assessed by iterative mapping using BOWTIE2 2.3.4.3 (38,39) and by whole genome alignment; closest relatives were identified through similarity searches, as performed above using MAFFT v7.450 (40,41). Open reading frames (ORFs) were identified and annotated using ORF finder in Geneious Prime 2021.0.3 (42). Only one contig displayed a complete coronavirus coding sequence (CDS). Its corresponding taxa was temporarily named “mute swan-associated gammacoronavirus”.

### RT-PCR screening

The genetic diversity of the gammacoronaviruses circulating in our study habitat was assessed using a two-step RT-PCR assay targeting the nsp10-12 domain of the coronavirus polymerase gene, followed by Sanger sequencing. The screening was conducted on 964/1080 mute swan faecal samples and 671 faecal samples from other bird species using a two-dimensional pooling approach. Samples were pooled separately for mute swans and for non-mute swan birds, along the two axes of a 10×10 grid, resulting in 196 pools of mute swan samples and 139 pools of other bird species samples (**supplementary table 2**). RNA from each pool was extracted using MagMAX CORE Nucleic Acid Purification Kit (ThermoFisher Scientific) and KingFisher™ Flex Purification System (ThermoFisher Scientific) following manufacturer’s instructions. Reverse transcription was performed using Superscript III reverse transcriptase (ThermoFisher Scientific). cDNA was amplified using a specific primer pair designed using Primer-BLAST (NCBI), targeting 995 base pairs of the nsp10/12 peptide (Coronavirus_nsp10-12_F: 5’-ACGAAGCGCAATGTAATGCC-3’, Coronavirus_nsp10-12_R: 5’-AAGGGCAACATAACGCTCCA-3’). PCR was completed using Q5 Hot Start High Fidelity Polymerase (NEB) according to the manufacturer’s instructions, with 0.5 μM final concentration of each primer and 2.5 μl cDNA per 25 μl reaction. Amplification conditions were 98 °C for 30 seconds to activate the DNA polymerase, followed by 40 cycles of denaturation at 98 °C for 10 seconds, primer annealing at 62 °C for 15 seconds, and chain elongation at 72 °C for 45 seconds, followed by a final extension at 72 °C for 2 minutes. Amplification of 995 nt size bands was assessed through gel electrophoresis, using 1.5% agarose gels dyed with ethidium bromide. All positive amplicons were purified by adding 0.5 U of Exonuclease I (NEB) and 0.25 U of Shrimp Alkaline Phosphatase (NEB) to 20 μl of each PCR product, and adjusted to 30 μl using nuclease free water. The mixture was incubated at 37 °C for 30 minutes and 85 °C for 15 minutes to remove single-stranded DNA and free dNTPs. All purified PCR products were sequenced by Sanger sequencing. Chromatograms were checked for disparities between the forward and reverse reads, and consensus sequences for each amplicon were aligned to the mute swan-associated gammacoronavirus genome by MAFFT v7.450 using the E-INS-i algorithm to confirm their identity. Where Sanger sequences were generated from pooled samples, all resultant chromatograms were checked for the absence of minor and mixed peaks to ensure that sequences were not chimeric. Amplicons were also generated from samples obtained from individual birds where these had been included in a previous metagenomic study (30).

### Taxonomic attribution and phylogenetic analysis

A phylogenetic tree was estimated in order to place the 75 bird-associated nsp10/12 sequences we obtained from RT-PCR screening Sanger sequencing within the currently-known diversity of the *Gammacoronavirus* genus. All nucleotide sequences for the nsp10-12 domain of the 1ab protein belonging to the *Gammacoronavirus* genus were downloaded from GenBank on 7^th^ February 2021, comprising 1409 sequences. An initial alignment of all these sequences was performed by MAFFT using the FFT-NS-2 algorithm, refined by the G-INS-i algorithm (41,43), and was used to construct a neighbour-joining tree with 100 bootstrap replicates. To infer the putative number of coronavirus species represented by these sequences, we conducted a clustering analysis using the species demarcation threshold ratified by the ICTV: sequences were converted to amino acids, and clustering analysis was conducted on them using Uclust (44), with 90% amino acid identity used as a clustering threshold (45). All the 75 coronavirus sequences generated here formed a monophyletic clade supported by a bootstrap score >90%; these were extracted and realigned using the E-INS-i algorithm. That alignment was then used to estimate a maximum likelihood (ML) tree in PhyML 3.3 (46) under an HKY85 substitution model with gamma distributed rate variation among sites, estimation of the proportion of invariant sites, and unequal base frequencies (as chosen by jModelTest (47)) using 100 bootstrap replicates. The ML tree was mid-point rooted and visualised with FigTree 1.4 (http://tree.bio.ed.ac.uk/software/figtree/). This ML tree grouped all the sequences of a putative gammacoronavirus species, which we hereafter refer to as “*Waterbird gammacoronavirus 1*”. In order to estimate a minimum number of host species changes compatible with the virus phylogeny, we undertook a maximum likelihood ancestral character state analysis using TreeTime 0.8.5 (48). The phylogenetic tree used for this analysis contained 79 waterbird gammacoronavirus 1 sequences for which host information was available.

### Recombination analysis

Two complementary approaches were used to detect putative recombination events involving the two waterbird gammacoronavirus 1 complete coding sequences (i.e. the mute swan-associated gammacoronavirus genome detected here, and a Canada goose coronavirus (NC_046965)) and all other bird-infecting gammacoronaviruses (24 complete genomes on GenBank, as of 11^th^ January 2021). Prior to recombination analysis, the 26 genomes were aligned by MAFFT v7.450 using the FFT-NS-2 algorithm, and the alignment was refined using the G-INS-i algorithm.

In the first approach, the alignment was analysed for recombination using the RDP (49), GENECONV (50), Chimaera (51), MaxChi (52), BootScan (53), SiScan (54), Phylpro (55) and 3Seq (56) methods implemented in RDP4 v101 (57), with a window size of 400nt and a step size of 200nt. Multiple comparison test adjustment of p-values was performed using Bonferroni correction (p-value significance threshold ≤ 0.01). For the RDP method, internal references were used. We considered as significant only those recombination events detected by >3 methods. Potential recombination events suggested by RDP4 were investigated further by similarity plot and bootscan analyses (1000 bootstrap replicates) using SimPlot 3.5.1. (58).

In the second approach, maximum likelihood trees for four subgenomic regions (defined by putative recombination breakpoint locations) were estimated in order to investigate the phylogenetic origin of parental genome regions. The breakpoint locations were based on pairwise alignment of the two waterbird gammacoronavirus 1 genomes. Trees were estimated by RAxML (59) under the GTR + GAMMA nucleotide model (as chosen by jModelTest) using 1000 bootstrap replicates. The reference genome of beluga whale coronavirus SW1 (NC_010646) was used as an outgroup. Trees were formatted using FigTree. A list of all the gammacoronavirus genomes used in the recombination analyses, and their associated metadata, is provided in **supplementary table 3**.

### Prevalence modelling

A 2-level Bayesian hierarchical logistic regression model was developed to estimate both sample-level and pool-level prevalence of waterbird gammacoronavirus 1 within our pooled RT-PCR testing regime for mute swan faecal samples. The detection status of each of the 964 individual samples was assumed to follow a Bernoulli distribution with parameter *sample*_*i*_, the probability of sample *i* being positive. *sample*_*i*_ was expressed as a logistic regression with temporal and biological factors as covariates, namely meteorological seasons, epidemiological years (running 1^st^ June – 31^st^ May) and swan age-classes (cygnets <1 year old, juveniles 1–3 years old, and adults >3 years old; (60); **supplementary table 4**). Additional information on handling of samples from birds with no known age is provided in the supplementary material. Regression coefficients were drawn from weakly informative prior distributions *Normal*(0,100). The RT-PCR status of a sampling pool *j* was then also assumed to follow a Bernoulli distribution with probability of success expressed as:

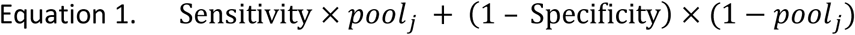

*Pool*_*j*_ was equal to 1 if at least one of the samples included in pool *j* was positive, and 0 otherwise. Sensitivity was defined as the probability of a pool to test positive if it included at least one positive sample, and the specificity as the probability of a pool to test negative if all samples in this pool would test negative. The inclusion of sensitivity and specificity allowed us to account for possible discrepancies between the observed and unobserved PCR statuses of a pool and its samples, respectively. As such discrepancies were expected to be rare, high sensitivity and specificity were assumed, and very informative prior distributions, Beta(400,1), were used.

The model was run using a Markov chain Monte Carlo simulation in JAGS 4.3.1 (61) and R.4.2.2 (62). A burn-in period of 10,000 iterations was used followed by a further 40,000 iterations which enabled full convergence in all parameters (Gelman and Rubin statistic <1.01; effective sample sizes >1,000). For all parameters, we report median and highest density intervals (HDI). All possible combinations of regression covariates were tested, and the model with the lowest deviance information criterion (DIC) was considered as the best supported model if the difference in DIC value was >5 from the next best performing model (**supplementary table 5**). A posterior predictive check was performed to assess models’ goodness of fit. Parameter values were sampled from the joint posterior, the PCR status of each sample and pool was simulated, and the resulting number of positive pools was compared to the observations. The posterior predictive sample-level prevalence was calculated by taking the inverse logit transformation of the corresponding regression coefficients sampled from the joint posterior distribution.

## 5. Results

### Taxonomic attribution and genome organisation of the new gammacoronavirus isolate

A 28,467 nt genome that encodes all the proteins of a gammacoronavirus was assembled from one mute swan sample (**Figure 1a**). The genome was briefly mentioned in Hill et al (2022) but is characterised and investigated in detail here. Whilst the terminal ends of both UTRs are missing, coverage across the genome is high (95% of the genome is covered by

**Figure 1:**
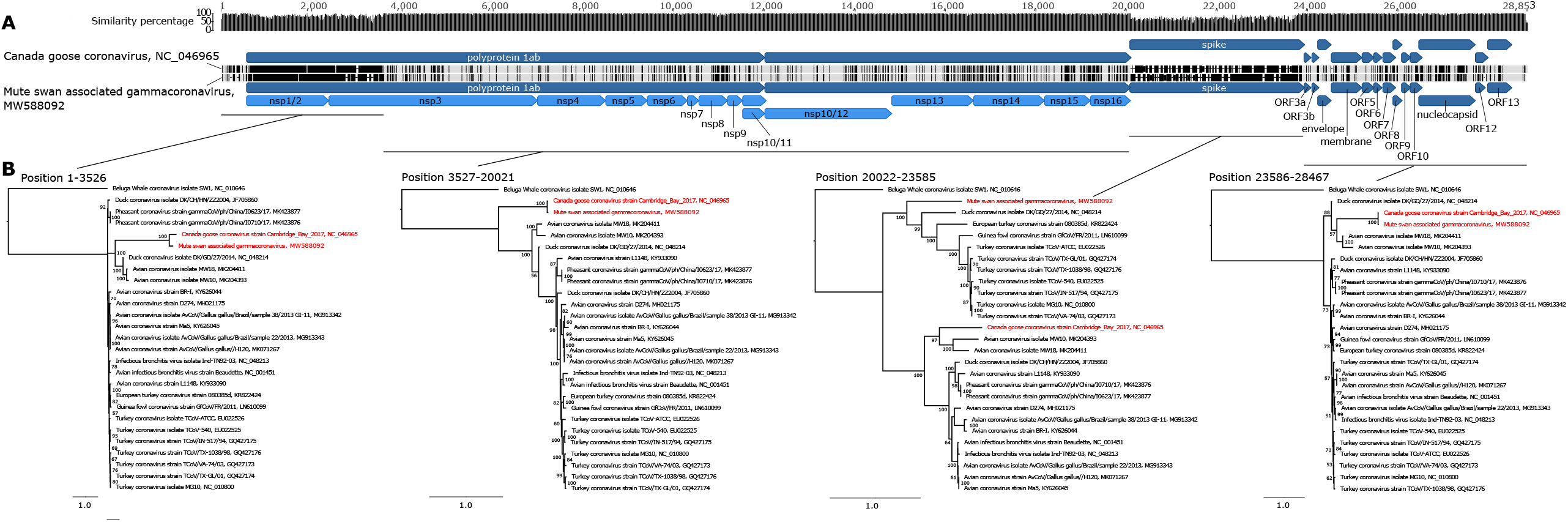
(A) Organisation of the two complete genomes of waterbird gammacoronavirus 1. Mismatches between the two aligned sequences are highlighted in black and a plot of sequence similarity along the genome is shown at the top of the panel. Dark blue: ORFs; light blue: peptides. (B) Maximum likelihood phylogenetic trees of 26 bird gammacoronavirus genomes were estimated for four subgenomic regions. Bootstrap scores (1000 replicates) are indicated at each node. Scale bars indicate nucleotide substitutions per site. Beluga whale coronavirus isolate SW1 (NC_010646) was used to root the trees. Genomes belonging to the *Waterbird gammacoronavirus 1* species are shown in red.

>1000 reads, with an average depth of 1907 reads, standard deviation=580) for a total of 395,069 mapped reads. This genome is named here ‘mute swan-associated gammacoronavirus’ (abbreviated as MSG).

MSG’s closest relative, in terms of genome sequence similarity, is the Canada goose coronavirus Cambridge_Bay_2017 (NC 046965, subsequently abbreviated as CGG; **Figure 1, supplementary table 6**). The CGG genome is the sole described member of the *Brangacovirus* sub-genus of the Gammacoronavirus genus, and belongs to the *Goose coronavirus CB17* species. The CGG genome was isolated from *Branta canadensis* (Canada goose) which, like the mute swan, belongs to the Anatidae waterbird family (63). According to the ninth report of the International Committee on Taxonomy of Viruses (ICTV), coronaviruses that share >90% amino acid (aa) sequence identity in seven conserved replicase domains (i.e. nsp3, nsp5, nsp12, nsp13, nsp14, nsp15, and nsp16 located in the 1ab gene) are considered to belong to the same species. This 90% identity threshold serves as the sole species demarcation criterion (45). As MSG and CGG share 94.8% amino acid identity across their entire 1ab protein (**supplementary table 6**), they thus represent two isolates of the same species. Since our finding demonstrates that the host range of this virus species is not limited to Canada geese, we propose that the species name should be updated to reflect the wider host range, and here refer to the species as “*Waterbird gammacoronavirus 1*”.

The number and lengths of open-reading-frames of MSG are conserved compared to those of CGG. The two genomes share 88.2% nucleotide identity overall (93.6% in the 1ab protein gene, 98.0% in the envelope protein gene, 96.9% in the membrane protein gene and 95.2% in the nucleocapsid protein gene; **supplementary table 6**). However, the two genomes have substantially different sequences in the spike protein gene (only 49.1% nucleotide identity) and at the beginning of the gene encoding the 1ab protein (**Figure 2a**). This suggests that potential recombination could have occurred between one or both waterbird gammacoronavirus 1 genomes and other coronaviruses during their evolutionary history.

**Figure 2:**
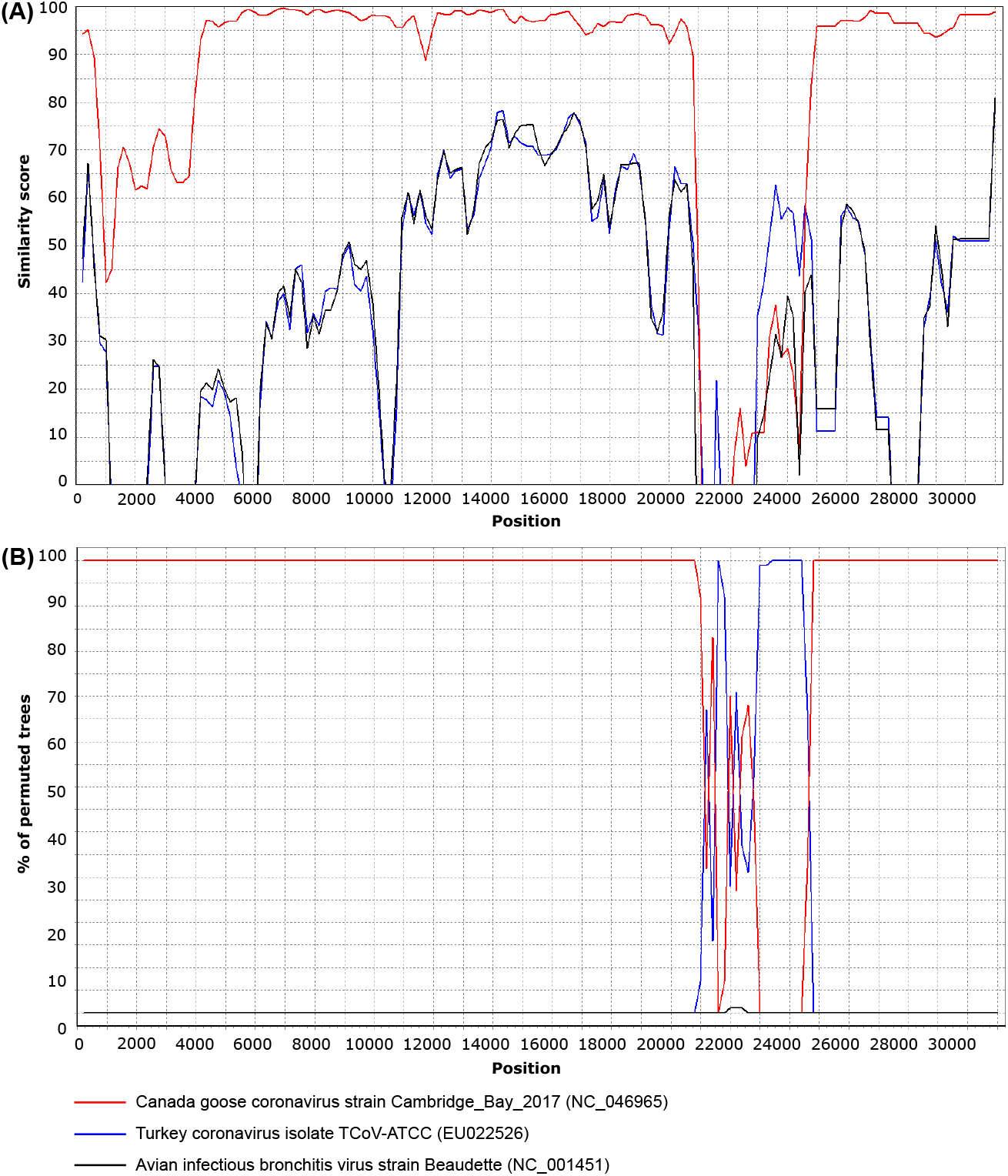
Detection of recombination in the genome of waterbird gammacoronavirus 1, isolate mute swan-associated gammacoronavirus, by (A) similarity plot and (B) bootscan analysis. Analysis was conducted with Simplot version 3.5.1 (window size: 400 bp; step size: 200 bp). Mute swan-associated gammacoronavirus (MW588092) was used as a query sequence and compared with the genome sequences of Canada goose coronavirus (NC_046965, red), turkey coronavirus isolate TCoV-ATCC (EU022526, blue) and avian infectious bronchitis virus strain Beaudette (NC_001451, black).

### Recombination event detection in one waterbird gammacoronavirus 1 isolate

The two waterbird gammacoronavirus 1 genomes were aligned with their closest relatives, i.e. all 24 complete genomes belonging to the *Igacovirus* sub-genus available from the GenBank database. This alignment was used to search for recombination events using eight recombination detection methods implemented in RDP4, combined with analysis of phylogenetic trees estimated from four subgenomic regions defined by the putative recombination breakpoints. Potential recombination events were investigated further by similarity plot and bootscan analyses.

The MSG and CGG genomes clustered together in all subgenomic phylogenetic trees, except for the spike protein phylogeny, in which these two lineages are separated (**Figure 1**). This suggests that either the spike protein gene of CGG or of MSG likely originated from a recombination event. In all phylogenies except that of the spike protein gene, MSG and CGG share a common ancestor with avian coronavirus isolates MW18 (MK204411) and MW10 (MK204393), implying that the at least one spike protein has become inserted into a parental genome representing that main evolutionary history. In the spike protein gene phylogeny, MSG does not cluster with CGG, MW18 and MW10, implying that MSG is the recombinant virus. This putative recombination event was confirmed by six recombination detection methods (BootScan, RDP, Chimaera, MaxChi, SiScan, and Phylpro, p-values <0.001). MSG could have originated from a recombination event between two parental viruses: (i) a virus with regions of the genome outside the spike protein that are closely related to CGG (averaging 94.5% nucleotide identity) and (ii) an undescribed gammacoronavirus whose spike protein is likely more closely related to viruses in the clade containing turkey coronavirus (EU022526) with which MSG shares 57.1% nucleotide identity (64) (**Figure 2, supplementary table 6**). All the aforementioned clades in phylogenetic trees are supported by >95% bootstrap support, therefore we conclude that different phylogenetic placements of the MSG lineage constitute good evidence of a recombination event during the history of this virus.

Recombination breakpoints – i.e. the positions defining genomic regions with different ancestral histories – were estimated to be located at positions 19,794 (99% CI: 19,622-20,340) and 23,436 (99% CI: 23,393-23,507) of the MSG genome, as visualised by a similarity plot (**Figure 2**). These positions are very close to the first and last nucleotide positions of the MSG spike protein (20,012 and 23,593 respectively) indicating that almost the entire spike protein gene originated from this recombination event. This result is congruent with our current knowledge of recombination hotspots in coronaviruses. We undertook an informal literature survey of 25 research papers on animal coronavirus recombination (**supplementary table 7**) and found that the spike protein gene is the most reported hotspot of recombination events in orthocoronaviruses (reported in 21/25 papers), and that the area immediately upstream of the spike protein gene shows the highest number of recombination breakpoints (**supplementary table 7**).

In addition to low sequence similarity between the two isolates of the waterbird gammacoronavirus 1 genomes (CGG and MSG) in their spike protein gene, we observed low sequence similarity between the two isolates at the 5’ end of the 1ab protein gene. These do not appear to have arisen through recombination. The recombination detection methods implemented in RDP4 reported no recombination in this region and the phylogenetic topology of the region matches that of the other non-spike genome regions (**Figure 2**). It is possible that the observed divergence was caused by either MSG or CGG recombining with a non-described gammacoronavirus that was a close relative of MSG and CGG. This is feasible because of our very limited knowledge of the genomic diversity of gammacoronaviruses, especially those that occur in wildlife. Alternatively, MSG and CGG could evolve faster in this genome region than in other genome regions (excluding the spike gene), as illustrated by the longer terminal branches leading to MSG and CGG in the 1ab gene phylogeny (**Figure 1b**).

### Genetic diversity of waterbird gammacoronavirus 1 in wild waterbirds

The genetic diversity of the waterbird gammacoronavirus 1 circulating in our study habitat was investigated by Sanger sequencing of RT-PCR amplified fragments of the nsp10/12 domain. The nsp10/12 domain was chosen because (i) it is the most conserved genomic region of coronaviruses, and (ii) it was investigated in previous studies that have attempted to explore coronavirus diversity (65–67).

A 995 nt fragment of the nsp10/12 peptide of the viral polymerase gene was successfully sequenced from 8 samples for which gammacoronavirus sequences were detected in high-throughput sequencing reads, and from 67 pools of faecal material from swans and other bird species, representing 75 non-chimeric sequences in total. Phylogenetic analysis of these nsp10/12 sequences indicates that these 75 sequences fall inside a monophyletic clade that also comprises 20 sequences from GenBank, and which share >90% nucleotide identity (**Figure 3**). All these partial genomes therefore likely belong to the *Waterbird gammacoronavirus 1* species, according to the ICTV coronavirus species criteria (45).

**Figure 3:**
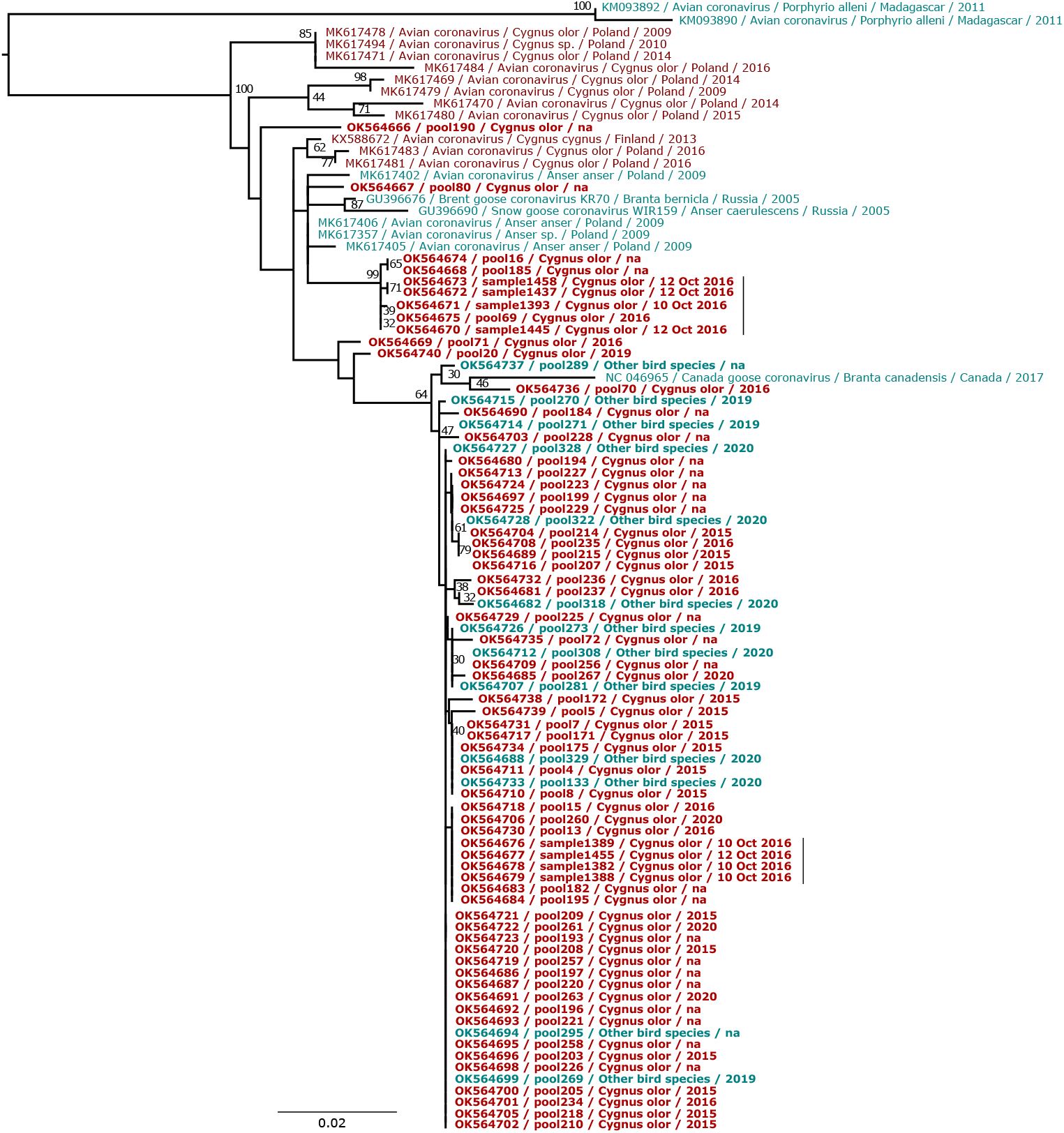
Maximum likelihood phylogenetic tree based on the nsp10-12 domain of the 1ab protein gene of 95 waterbird gammacoronavirus 1 sequences. The tree is mid-point rooted. Bootstrap values (100 replicates) higher than 30% are indicated at each node. Scale bar corresponds to nucleotide substitutions per site. The sequences from swans (*Cygnus spp*.) are in red, and the sequences from other bird species are in blue. The sequences obtained from our samples are in bold.

The sequences sampled from mute swans are diverse (89.7% to 100% nucleotide identity in the sequenced nsp10/12 region) and distributed throughout the phylogeny (**Figure 3**).

Notably, sequences isolated from samples collected at our study site on three consecutive days in October 2016 cluster in two separate clades (samples 1437, 1393, 1445, 1458; and 1382, 1388, 1389, 1455) (**Figure 3**). This indicates that multiple lineages of waterbird gammacoronavirus 1 were co-circulating within the bird community at our study site at that time.

Eighty of the waterbird gammacoronavirus 1 sequences are known to have been isolated from geese (*Anser sp*. and *Branta spp*.) or swans (*Cygnus spp*.) (**Figure 3**). These birds are taxonomically related and belong to the Anserinae sub-family of the Anatidae family; they also share similar habitats. The two remaining waterbird gammacoronavirus 1 sequences are more distantly related, and were found in Madagascar in the waterbird *Porphyrio alleni* (Allen’s gallinule; formerly *Porphyrula alleni*), which belongs to the Rallidae. Given the intermingled positions of waterbird gammacoronavirus 1 sequences from geese and swans in the phylogenetic tree (**Figure 3**), cross-species transmission of this virus between these birds is likely common. Maximum likelihood ancestral character state reconstruction indicates that a minimum of one host state change could separate swan-infecting and Rallidae-infecting virus lineages, and that at least two cross species transmission events from swans to geese occurred during the evolutionary history of this virus (**supplementary figure 1**). The frequency of cross-species transmission of waterbird gammacoronavirus 1 among geese and swans (or other waterbird species) in waterbird communities should be investigated further, especially since the virus appears to have a wide geographical distribution (encompassing at least Europe, Canada and Madagascar) (**Figure 3**) (63).

### Seasonality of waterbird gammacoronavirus 1 detection in wild waterbirds

A 2-level Bayesian hierarchical logistic regression model was developed to estimate both sample-level and pool-level prevalence of waterbird gammacoronavirus 1 within mute swan samples studied using our pooled RT-PCR testing regime. The best supported model for the 964 mute swan samples contained only “season” as an explanatory variable, and was able to produce a predictive number of positive sampling pools (median: n = 89, 95% HDI: n = 67–110) that was consistent with the observed value (n = 74) (**supplementary figure 2**). This model suggested that sample-level waterbird gammacoronavirus 1 prevalence varied across seasons (**Figure 4**), with lowest posterior predicted prevalence found in summer (1.1%, 95% HDI: 0.3–1.8%) and highest in spring (17.7%, 95% HDI: 10.7–25.2%) (**supplementary table 8**). Sample-level prevalence was not found to vary with epidemiological year or swan age classes.

**Figure 4:**
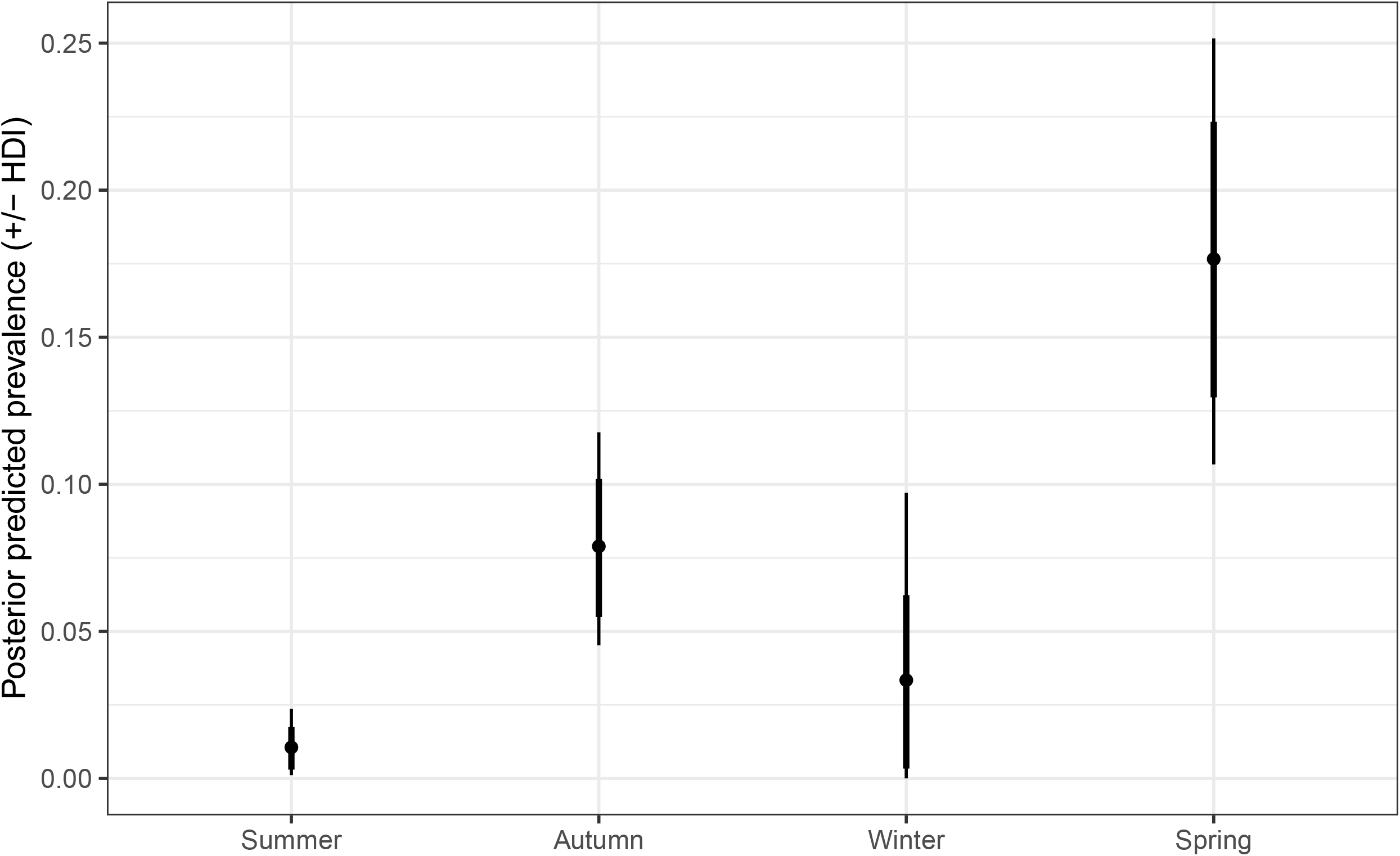
Posterior predicted sample-level prevalence for waterbird gammacoronavirus 1 across meteorological seasons. Error bars indicate 95% (thin bar) and 80% (thick bar) highest density intervals, with the central point indicating the median value.

## 6. Discussion

The importance of understanding coronavirus cross-species transmission is exemplified by the emergence in humans of three zoonotic betacoronaviruses during the last twenty years. Our work is predicated on the notion that predicting risks of future emerging viruses requires an understanding of the factors that explain virus transmission dynamics, genetic diversity, and evolution in wildlife reservoir populations and in food production systems.

Here, we characterised the presence and genetic diversity of a waterbird gammacoronavirus across multiple years, in a complex natural habitat containing multiple interacting waterbird species. This virus species was first identified in Canada geese and its classification has been recently ratified by the ICTV under the name *Goose coronavirus CB17* (63,68). Since our findings show that the host range of this virus species also includes swans, we propose that the species name should be updated to reflect the wider host range, and we propose “*Waterbird gammacoronavirus 1*”. Importantly, viruses likely belonging to *Waterbird gammacoronavirus 1* have been isolated from distant geographical locations since 2011, including Eastern Europe, Madagascar and Canada, suggesting a wide and persistent distribution of the virus (**Figure 3**). Non-invasive sampling of waterbird gammacoronavirus 1, and related coronaviruses, may therefore provide a tractable model system for understanding the evolutionary and cross-species dynamics of cororaviruses in the wild.

Our modelling estimated that waterbird gammacoronavirus 1 prevalence varied substantially by season in mute swans, ranging from 1.1% in summer to a seasonal peak of 17.7% in spring, with an overall average across the year of 7.5%. This prevalence is within the range reported for other gammacoronaviruses in wild Anatidae populations, which range between 3% and 45% (**supplementary table 5**) (27). Peaks in coronavirus detection during spring and autumn correlate with the passage and arrival of thousands of migratory birds of different species at our study site, including brent goose (*Branta bernicla*), northern pintail (*Anas acuta*), Eurasian teal (*Anas crecca*), common pochard (*Aythya ferina*), Northern lapwing (*Vanellus vanellus*), and Eurasian coot (*Fulica atra*) (69). Migration could increase virus abundance through introduction of new viral lineages, and/or by increasing host abundance and density. This would be consistent with the observed concurrent introduction and circulation of several genetically diverse lineages (**Figure 3**). We note that several highly pathogenic avian influenza outbreaks have been introduced in our study population during the last decade, likely via migration of other species (70–72).

The host range of waterbird gammacoronavirus 1 is likely broad and includes at the very least swans and geese (belonging to the *Cygnus, Anser* and *Branta* genera, Anatidae family) and the Allen’s gallinule (*Porphyrio* genus, Rallidae family). Birds from these families share similar aquatic habitats, which we hypothesise (i) facilitates coronavirus cross-transmission between different waterbird species, and (ii) increases the probability of viral recombination events between distant coronavirus lineages (73). We detected a recombination event in a waterbird gammacoronavirus 1 genome, involving its entire spike protein gene.

Recombination of the spike protein gene seems to occur commonly in bird-infecting coronaviruses, and has been documented for infectious bronchitis virus and turkey coronavirus species. Interestingly, the spike protein gene of turkey coronavirus likely originated from a unknown wildbird-infecting coronavirus (74,75). Our RT-PCR screening method targeted a conserved portion of the RdRp gene in order to allow us to robustly classify coronavirus-positive samples at to the level of virus species (45), but did not allow us to fully investigate genomic diversity in other gene regions (27). The characterisation of additional complete and divergent coronavirus genomes, using unbiased sequencing approaches, would improve our understanding of gammacoronavirus genomic diversity (27) and our discovery of recombinant genomes.

Our review of 25 research papers on animal coronavirus recombination shows that the spike protein gene is the most commonly reported location of coronavirus recombination events (reported in 21/25 studies) (**supplementary table 7**). The spike protein mediates host cell attachment, virus and cell membrane fusion and entry into the host cell and hence determines viral tropism (76). Hence, recombination events involving the spike protein gene can facilitate coronaviral host range adaptability, and consequently could be an important explanatory mechanism of coronaviruses host jumps. Accordingly, several bird coronaviruses have been reported in several wild bird species sharing aquatic habitats, such as ducks, gulls and shorebirds (27). Consistent with a recent study on waterbirds that did not show a clear association between virome composition and bird species from different orders (64), these results suggest that in natural settings where large populations of multiple species that freely mix, feed and defecate into the same shared habitat (watersource), host ecology could select for viruses with broad host range.

The pathogenicity of wild birds coronaviruses, including the waterbird gammacoronavirus 1, is unclear, and warrants further research. Infection with the Canada goose coronavirus strain (NC_046965) was detected in birds during a mass die-off of wild geese; however this observation is correlative and the aetiology of this outbreak is unknown (63). Conversely the mute swans we sampled that were positive for this virus species showed no obvious symptoms of disease at or shortly after sampling, none of the swans were reported dead within seven weeks after sampling, and there were not concurrent large die-offs observed in other species at the study site. This suggests no or low virulence of this virus. Moreover, we observed a high prevalence and genetic diversity of waterbird gammacoronavirus 1 in our population; if this prevalence is similar elsewhere then waterbird gammacoronavirus 1 could be commonly detected by accident in a population that is suffering mortality due to alternative causes.

Our results highlight that our knowledge of gammacoronavirus diversity and ecology is scarce, and biased towards domesticated animals. Our phylogenetic analysis of all available nsp10-12 domains of the RdRp gene from gammacoronaviruses (n=1409; **supplementary figure 3**) confirms that wild waterbirds are the reservoir of highly diverse and divergent gammacoronaviruses. Most of these sequences belong to partial genomes, and thus are not considered by the ICTV. However, the diversity of these sequences suggests they may represent at least 5 new gammacoronavirus species, which would double the number of gammacoronaviruses in waterbirds (4 species are currently defined). Future studies should ideally both confirm this with whole genome sequencing, and investigate whether the epidemiology of these putative new viruses is similar to that reported here for waterbird gammacoronavirus 1.

Wild aquatic birds are known to carry infectious agents that circulate within their communities with no or few signs of disease, notably low pathogenic avian influenza viruses (25). It is clear that to better prevent or mitigate the impacts of emerging pathogens we urgently need to expand genomic surveillance beyond human populations and undertake careful longitudinal studies at the interface between humans and wild animals, at the interface between wild animals and livestock/poultry, and within wild populations themselves. Studying cross-species transmission in multi-host systems in wildlife, integrating host ecology and virus genomic data, has a great potential to inform us on how viruses cross species boundaries and circulate in different hosts. Such studies are particularly important to improve our understanding of the ecology of the viruses of high zoonotic potential, such as coronaviruses.

## 7. Author statements

### 7.2 Authors and contributors

S.F.: Conceptualisation, Methodology, Validation, Formal Analysis, Investigation, Data Curation, Writing-Original Draft Preparation, Writing-Review and Editing, Visualisation, Supervision, Funding

S.N.: Investigation, Validation, Writing-Review and Editing, Funding S.H.V.: Formal Analysis, Writing-Review and Editing

G.F.: Methodology, Formal Analysis, Writing-Review and Editing C.M.P.: Investigation

A.J.B.: Resources, Writing-Review and Editing, Supervision, Project Administration, Funding. O.G.P.: Conceptualisation, Methodology, Writing-Review and Editing, Supervision, Project Administration, Funding

S.C.H.: Conceptualisation, Methodology, Investigation, Formal Analysis, Writing-Review and Editing, Supervision, Project Administration, Funding

### 7.4 Ethical approval

Not applicable.

### 7.5 Consent for publication

Not applicable.

## 7.6 Acknowledgements

We thank Mrs C. Townsend for permission to study the Abbotsbury swans, and the staff at Abbotsbury Swannery for helping to collect the demographic data.

