## Supplementary materials for "Genetic diversity, recombination and cross-species transmission of a waterbird gammacoronavirus in the wild"

### Supplementary Material

**Supplementary figure 1**: Maximum likelihood phylogenetic tree based on the nsp10-12 domain of 79 waterbird gammacoronavirus 1 sequences upon which ancestral trait state reconstructions have been superimposed. Internal node colours indicate the parsimonious ancestral trait state reconstruction of the host genus, as shown in the key. The tree is mid-point rooted. Bootstrap values (1000 replicates) higher than 50% are indicated at each node. Scale bar corresponds to nucleotide substitutions.

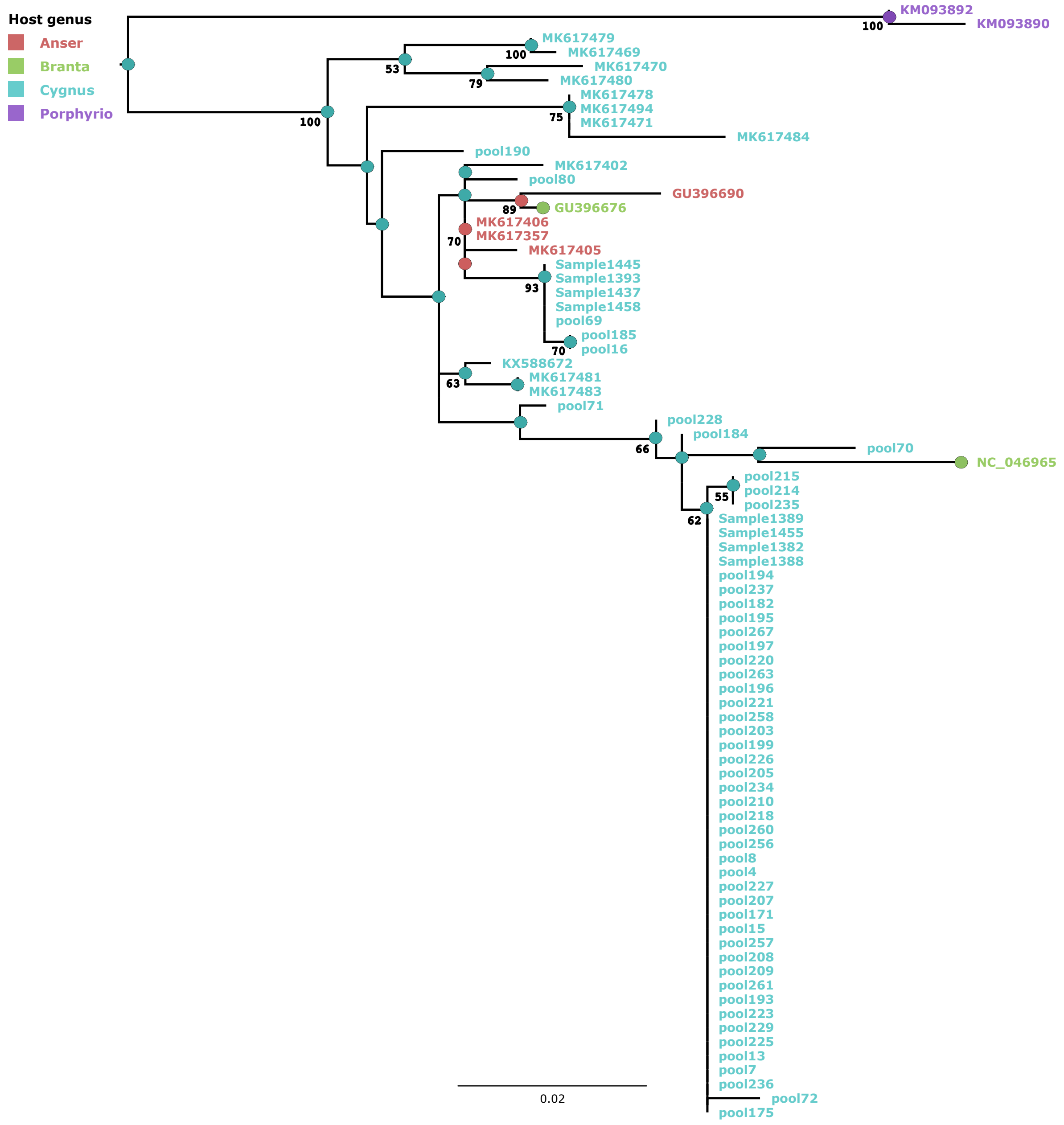

**Supplementary figure 2**: Posterior predictive distribution of positive RT-PCR pools. Vertical dashed red lines indicate 95% highest density intervals. Vertical solid red line indicates the observed positive number of pools.

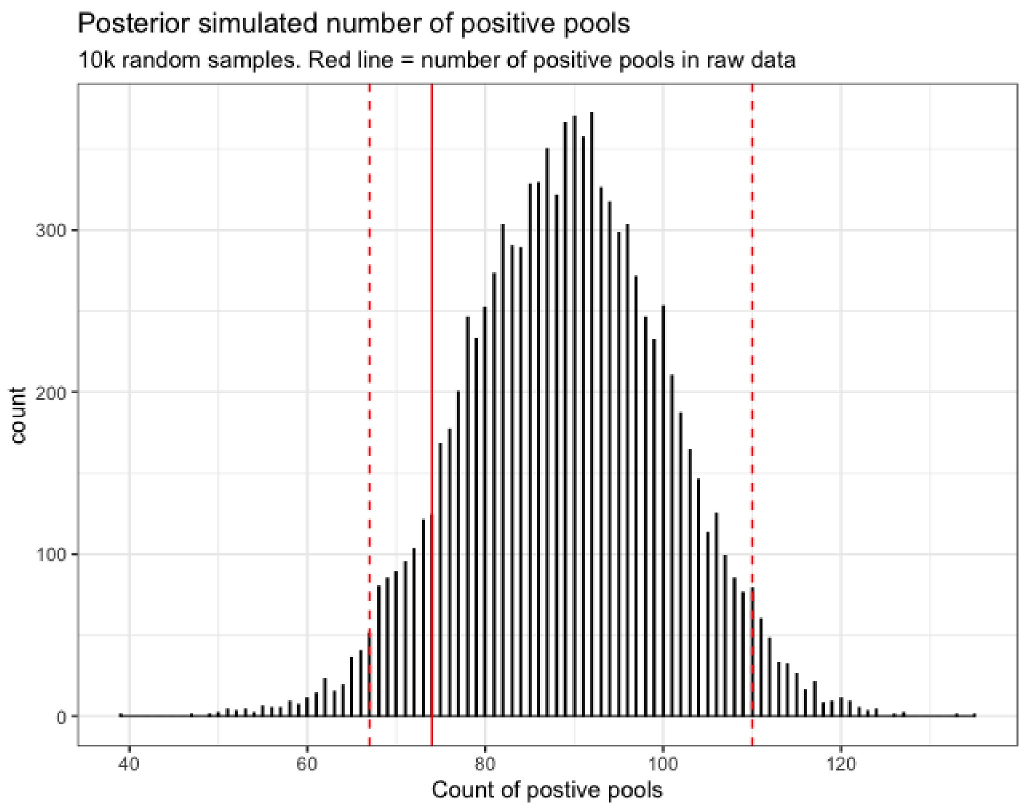

**Supplementary figure 3**: Maximum likelihood phylogenetic tree based on the nsp10-12 domain of the 1ab protein gene of 1429 gammacoronavirus sequences. Major bootstrap values (100 replicates) higher than 50% are indicated at each node. Scale bar corresponds to nucleotide substitutions per site. The sequences obtained from our samples are coloured in red. Hosts from which viral sequences were isolated are indicated by silhouettes. Representative species members are marked in bold. Blue circles indicate branches defining putative species clades according to ICTV recommendation.

**Supplementary table 1**: List of the 1751 faecal samples used in this study, and of their associated metadata.

**Supplementary table 2**: Grid plans of the processed pools. RT-PCR positive pools are marked in green.

**Supplementary table 3**: List of the 27 genomes of gammacoronaviruses, including 26 bird-infecting coronaviruses and a beluga whale-infecting coronavirus, that were used for phylogenetic and recombination analyses.

| GenBank accession number | Species name (ICTV) | Isolate name | Sub-genus | Collection date | Country | Isolation source (scientific name) | Isolation source (common name) |
|---|---|---|---|---|---|---|---|
| NC_046965 | Goose coronavirus CB17 | Canada goose coronavirus strain Cambridge_Bay_2017 | Brangacovirus | 2017 | Canada | *Branta canadensis* | Canada goose |
| MW588092 | Goose coronavirus CB17 | Mute swan associated gammacoronavirus Abbotsbury_2016 | Brangacovirus | 2016 | United Kingdom | *Cygnus olor* | mute swan |
| NC_048213 | Avian coronavirus | Infectious bronchitis virus isolate Ind-TN92-03 | Igacovirus | 2003 | India | *Gallus gallus* | chicken |
| NC_001451 | Avian coronavirus | Avian infectious bronchitis virus strain Beaudette | Igacovirus | | | *Gallus gallus* | chicken |
| KY626044 | Avian coronavirus | Avian coronavirus strain BR-I | Igacovirus | 2016 | Brazil | *Gallus gallus* | chicken |
| MG913342 | Avian coronavirus | Avian coronavirus isolate AvCoV/Gallus gallus/Brazil/sample 38/2013 GI-11 | Igacovirus | 2013 | Brazil | *Gallus gallus* | chicken |
| KY626045 | Avian coronavirus | Avian coronavirus strain Ma5 | Igacovirus | 2016 | Brazil | *Gallus gallus* | chicken |
| MK071267 | Avian coronavirus | Avian coronavirus strain AvCoV/Gallus gallus/H120 | Igacovirus | | Brazil | *Gallus gallus* | chicken |
| MG913343 | Avian coronavirus | Avian coronavirus isolate AvCoV/Gallus gallus/Brazil/sample 22/2013 | Igacovirus | 2013 | Brazil | *Gallus gallus* | chicken |
| MH021175 | Avian coronavirus | Avian coronavirus strain D274 | Igacovirus | 1979 | Netherlands | *Gallus gallus* | chicken |
| KY933090 | Avian coronavirus | Avian coronavirus strain L1148 | Igacovirus | | | *Gallus gallus* | chicken |
| JF705860 | Avian coronavirus | Duck coronavirus isolate DK/CH/HN/ZZ2004 | Igacovirus | 2004 | China | *Anas platyrhynchos* | mallard |
| MK423876 | Avian coronavirus | Pheasant coronavirus strain gammaCoV/ph/China/I0710/17 | Igacovirus | 2017 | China | *Phasianus colchicus* | pheasant |
| MK423877 | Avian coronavirus | Pheasant coronavirus strain gammaCoV/ph/China/I0623/17 | Igacovirus | 2017 | China | *Phasianus colchicus* | pheasant |
| NC_010800 | Avian coronavirus | Turkey coronavirus isolate MG10 | Igacovirus | | Canada | *Meleagris gallopavo* | turkey |
| GQ427173 | Avian coronavirus | Turkey coronavirus strain TCoV/VA-74/03 | Igacovirus | 2003 | USA | *Meleagris gallopavo* | turkey |
| GQ427175 | Avian coronavirus | Turkey coronavirus strain TCoV/IN-517/94, GQ427175 | Igacovirus | 1994 | USA | *Meleagris gallopavo* | turkey |
| EU022525 | Avian coronavirus | Turkey coronavirus isolate TCoV-540 | Igacovirus | | USA | *Meleagris gallopavo* | turkey |
| GQ427174 | Avian coronavirus | Turkey coronavirus strain TCoV/TX-GL/01 | Igacovirus | 2001 | USA | *Meleagris gallopavo* | turkey |
| GQ427176 | Avian coronavirus | Turkey coronavirus strain TCoV/TX-1038/98 | Igacovirus | 1998 | USA | *Meleagris gallopavo* | turkey |
| EU022526 | Avian coronavirus | Turkey coronavirus isolate TCoV-ATCC | Igacovirus | | USA | *Meleagris gallopavo* | turkey |
| LN610099 | Avian coronavirus | Guinea fowl coronavirus strain GfCoV/FR/2011 | Igacovirus | 2011 | France | *Numididae* | guinea fowl |
| KR822424 | Avian coronavirus | European turkey coronavirus strain 080385d | Igacovirus | 2008 | France | *Meleagris gallopavo* | turkey |
| NC_048214 | Duck coronavirus 2714 | Duck coronavirus isolate DK/GD/27/2014 | Igacovirus | 2014 | China | *Anas platyrhynchos* | mallard |
| MK204393 | UNCLASSIFIED | Avian coronavirus isolate MW10 | Igacovirus | 2017 | Australia | *Anas gracilis* | grey teal |
| MK204411 | UNCLASSIFIED | Avian coronavirus isolate MW18 | Igacovirus | 2017 | Australia | *Tadorna tadornoides* | Australian shelduck |
| NC_010646 | Beluga whale coronavirus SW1 | Beluga Whale coronavirus isolate SW1 | Cegacovirus | | USA | *Delphinapterus leucas* | beluga whale |

**Supplementary table 4**: Sample sizes for prevalence modelling covariate groupings. Age information was missing for 5 individuals. We chose not to model these under an age-class 'Unknown' due to small sample size of this group and the risk of overfitting. To avoid removing these sample from our analysis and hence biasing pool-level positivity, the age-class of these individuals was simulated within the models that contained age as a covariate. To simulate age for the 5 unknown samples, seasonal age proportions of juveniles (excluding cygnets) for the remaining samples were calculated. Cygnets were removed from this as the 5 unaged individuals could be confirmed to be >1 years of age, but could not be definitively categorised as juvenile or adult. The simulated probability of juvenile age status of a sample *k* ($sim\_Age\_J_k$) was then assumed to follow a Bernoulli distribution with probability of success expressed as the relevant seasonal age proportion of juveniles for that sample. The Juvenile age-class status (0 or 1) of a sample *k* could then be expressed as;

Supplementary Materials Equation 1. $missing\_Age_k * sim\_Age\_J_k + (1-missing\_Age_k) * Age\_J_k$

where *missing_Age* was a vector of 0 or 1, where a value of 1 indicated age-class information was missing for that sample and parameter *Age_J* was a vector of 0 or 1 where a value of 1 indicated a known age of juvenile for that sample.

| | n (%) |
|---|---|
| All | 964 (100%) |
| Season | |
| Summer | 389 (40.4%) |
| Autumn | 310 (32.2%) |
| Winter | 100 (10.4%) |
| Spring | 165 (17.1%) |
| Epidemiological year | |
| 2014–2015 | 57 (5.9%) |
| 2015–2016 | 366 (38.0%) |
| 2016–2017 | 223 (23.1%) |
| 2017–2018 | 24 (2.5%) |
| 2019–2020 | 192 (19.9%) |
| 2020–2021 | 102 (10.6%) |
| Age | |
| Adult | 380 (39.4%) |
| Cygnet | 280 (29.0%) |
| Juvenile | 299 (31.0%) |

| | |
|---|---|
| Unknown | 5 (0.5%) |

**Supplementary table 5:** DIC comparisons for tested prevalence models

| Variable included in model? | | | DIC value |
|---|---|---|---|
| Season | Age | Epidemiological year | |
| ✓ | X | X | 239.1 |
| ✓ | ✓ | X | 250.2 |
| ✓ | X | ✓ | 258.5 |
| ✓ | ✓ | ✓ | 260.6 |
| X | X | X | 286.3 |
| X | X | ✓ | 291.0 |
| X | ✓ | X | 298.6 |
| X | ✓ | ✓ | 305.8 |

**Supplementary table 6**: Coding region and similarity of the waterbird gammacoronavirus 1 isolated from mute swans (*Cygnus olor*) to its closest relatives.

| ORF | Location (nt) | Length (nt) | Length (aa) | Closest relative isolate, GenBank accession number | | Percentage of amino acid identity | Percentage of nucleotide identity |
|---|---|---|---|---|---|---|---|
| polyprotein 1a | 539 - 11,944 | 11436 | 3825 | Waterbird gammacoronavirus 1, isolate Canada_goose/Cambridge_Bay/2017 | NC_046965 | 91.0% | 91.0% |
| polyprotein 1b | 11,944 - 20,028 | 8058 | 2685 | | | 99.3% | 97.2% |
| spike protein | 20,012 - 23,593 | 3582 | 1233 | Avian coronavirus, isolate Guinea fowl coronavirus GfCoV/FR/2011 | LN610099 | 57.6% | 60.6% |
| ORF3a | 23,586 - 23,747 | 162 | 53 | Waterbird gammacoronavirus 1, isolate Canada_goose/Cambridge_Bay/2017 | NC_046965 | 92.5% | 95.7% |
| ORF3b | 23,756 - 23,923 | 168 | 55 | | | 94.5% | 97.6% |
| envelope protein | 23,883 - 24,185 | 303 | 100 | | | 99.0% | 98.0% |
| membrane protein | 24,182 - 24,865 | 684 | 227 | | | 98.2% | 96.9% |
| ORF5 | 24,865 - 25,131 | 267 | 88 | | | 96.6% | 98.9% |

| | | | | | | | |
|---|---|---|---|---|---|---|---|
| ORF6 | 25,128 - 25,319 | 192 | 63 | | | 93.7% | 95.3% |
| ORF7 | 25,327 - 25,605 | 279 | 92 | | | 95.7% | 97.0% |
| ORF8 | 25,547 - 25,756 | 210 | 69 | | | 98.6% | 95.2% |
| ORF9 | 25,740 - 25,940 | 201 | 66 | | | 95.5% | 96.0% |
| ORF10 | 25,937 - 26,194 | 258 | 85 | | | 92.9% | 94.6% |
| nucleocapsid protein | 26,122 - 27,366 | 1245 | 414 | | | 95.9% | 95.2% |
| ORF12 | 27,371 - 27,592 | 222 | 73 | | | 87.7% | 95.5% |
| ORF13 | 27,640 - 28,182 | 543 | 180 | | | 95.6% | 98.3% |

**Supplementary table 7**: Literature review of research papers on coronavirus recombination in animals. The search was conducted on PubMed on 15th March 2021 with the keywords coronavirus AND recombination AND (animal OR bird). All the relevant papers were downloaded and processed, resulting in a table summarising the results of 25 research papers.

**List of the references used for the creation of supplementary table 7**

Gammacoronaviruses: (1)(2)(3)(4)(5)

Deltacoronaviruses: (6)(7)(8)(9)

Alphacoronaviruses: (10)(11)(12)(13)(14)(15)(16)(17)(18)(19)

Betacoronaviruses: (20)(21)(22)(23)(24)(25)


1. Jia W, Karaca K, Parrish CR, Naqi SA. A novel variant of avian infectious bronchitis virus resulting from recombination among three different strains. Arch Virol. 1995;140:259–71. doi: 10.1007/BF01309861
2. Lee CW, Jackwood MW. Evidence of genetic diversity generated by recombination among avian coronavirus IBV. Arch Virol. 2000;145(10):2135–48. doi: 10.1007/s007050070044
3. Lee CW, Jackwood MW. Spike gene analysis of the DE072 strain of infectious bronchitis virus: Origin and evolution. Virus Genes. 2001;22(1):85–91. doi: 10.1023/A:1008138520451
4. Thor SW, Hilt DA, Kissinger JC, Paterson AH, Jackwood MW. Recombination in avian gamma-coronavirus infectious bronchitis virus. Viruses. 2011;3(9):1777–99. doi: 10.3390/v3091777
5. Liu S, Xu Q, Han Z, Liu X, Li H, Guo H, et al. Origin and characteristics of the recombinant novel avian infectious bronchitis coronavirus isolate ck/CH/LJL/111054. Infect Genet Evol. 2014;23(April):189–95. doi: 10.1016/j.meegid.2014.02.015
6. Woo PCY, Lau SKP, Lam CSF, Lau CCY, Tsang AKL, Lau JHN, et al. Discovery of Seven Novel Mammalian and Avian Coronaviruses in the Genus Deltacoronavirus Supports Bat Coronaviruses as the Gene Source of Alphacoronavirus and Betacoronavirus and Avian Coronaviruses as the Gene Source of Gammacoronavirus and Deltacoronavi. J Virol. 2012;86(7):3995–4008. doi: 10.1128/jvi.06540-11
7. Lau SKP, Wong EYM, Tsang C-C, Ahmed SS, Au-Yeung RKH, Yuen K-Y, et al. Discovery and Sequence Analysis of Four Deltacoronaviruses from Birds in the Middle East Reveal Interspecies Jumping with Recombination as a Potential

Mechanism for Avian-to-Avian and Avian-to-Mammalian Transmission. J Virol. 2018;92(15):1–18. doi: 10.1128/jvi.00265-18

8. He WT, Ji X, He W, Dellicour S, Wang S, Li G, et al. Genomic epidemiology, evolution, and transmission dynamics of porcine deltacoronavirus. Mol Biol Evol. 2020;37(9):2641–54. doi: 10.1093/molbev/msaa117
9. Wang Q, Zhou ZJ, You Z, Wu DY, Liu SJ, Zhang WL, et al. Epidemiology and evolution of novel deltacoronaviruses in birds in central China. Transbound Emerg Dis. 2021;(February):1–13. doi: 10.1111/tbed.14029
10. Lau SKP, Woo PCY, Li KSM, Huang Y, Wang M, Lam CSF, et al. Complete genome sequence of bat coronavirus HKU2 from Chinese horseshoe bats revealed a much smaller spike gene with a different evolutionary lineage from the rest of the genome. Virology. 2007;367(2):428–39. doi: 10.1016/j.virol.2007.06.009
11. Wang W, Lin XD, Guo WP, Zhou RH, Wang MR, Wang CQ, et al. Discovery, diversity and evolution of novel coronaviruses sampled from rodents in China. Virology. 2015;474:19–27. doi: 10.1016/j.virol.2014.10.017
12. Lamers MM, Smits SL, Hundie GB, Provacia LB, Koopmans M, Osterhaus ADME, et al. Naturally occurring recombination in ferret coronaviruses revealed by complete genome characterization. J Gen Virol. 2016;97(9):2180–6. doi: 10.1099/jgv.0.000520
13. Jarvis MC, Lam HC, Zhang Y, Wang L, Hesse RA, Hause BM, et al. Genomic and evolutionary inferences between American and global strains of porcine epidemic diarrhea virus. Prev Vet Med. 2016;123:175–84. doi: 10.1016/j.prevetmed.2015.10.020
14. Tao Y, Shi M, Chommanard C, Queen K, Zhang J. Surveillance of Bat Coronaviruses in Kenya Identifies Relatives of Human Coronaviruses NL63 and 229E and Their Recombination History. J Virol. 2017;91(5):1–16. doi: 10.1128/JVI.01953-16
15. Wang W, Lin X-D, Liao Y, Guan X-Q, Guo W-P, Xing J-G, et al. Discovery of a Highly Divergent Coronavirus in the Asian House Shrew from China Illuminates the Origin of the Alphacoronaviruses. J Virol. 2017;91(17):1–15. doi: 10.1128/jvi.00764-17
16. Chen F, Knutson TP, Rossow S, Saif LJ, Marthaler DG. Decline of transmissible gastroenteritis virus and its complex evolutionary relationship with porcine respiratory coronavirus in the United States. Sci Rep. 2019;9(1):3953. doi: 10.1038/s41598-019-40564-z
17. Han Y, Du J, Su H, Zhang J, Zhu G, Zhang S, et al. Identification of diverse bat

alphacoronaviruses and betacoronaviruses in china provides new insights into the evolution and origin of coronavirus-related diseases. Front Microbiol. 2019;10(AUG). doi: 10.3389/fmicb.2019.01900

**Supplementary table 8:** Posterior predicted sample-level proportional prevalence for waterbird gammacoronavirus 1 across meteorological seasons.

| Season | Prevalence (proportion) | HDI (80%) | HDI (95%) |
|---|---|---|---|
| Summer | 0.011 | 0.003–0.018 | 0.001–0.024 |
| Autumn | 0.079 | 0.055–0.102 | 0.045–0.118 |
| Winter | 0.033 | 0.003–0.062 | 0.00002–0.097 |
| Spring | 0.177 | 0.130–0.223 | 0.107–0.252 |
